# Quantitative trait locus mapping identifies *Col4a6* as a novel regulator of striatal dopamine level and axonal branching in mice

**DOI:** 10.1101/2020.06.28.176206

**Authors:** Mélanie H. Thomas, Yujuan Gui, Pierre Garcia, Mona Karout, Christian Jaeger, Zdenka Hodak, Alessandro Michelucci, Heike Kollmus, Arthur Centeno, Klaus Schughart, Rudi Balling, Michel Mittelbronn, Joseph H. Nadeau, Robert W. Williams, Thomas Sauter, Lasse Sinkkonen, Manuel Buttini

## Abstract

The features of dopaminergic neurons (DAns) of nigrostriatal circuitry are orchestrated by a multitude of yet unknown factors, many of them genetic. Genetic variation between individuals at baseline can lead to differential susceptibility to and severity of diseases. As decline of DAns, a characteristic of Parkinson’s disease, heralds a significant decrease in dopamine level, measuring dopamine can reflect the integrity of DAns. To identify novel genetic regulators of the integrity of DAns, we used the Collaborative Cross (CC) mouse strains as model system to search for quantitative trait loci (QTLs) related to dopamine levels in the dorsal striatum. The dopamine levels in dorsal striatum varied greatly in the eight CC founder strains, and the differences were inheritable in 32 derived CC strains. QTL mapping in these CC strains identified a QTL associated with dopamine level on chromosome X containing 393 genes. RNA-seq analysis of the ventral midbrain of two of the founder strains with large striatal dopamine difference (C57BL/6J and A/J) revealed 24 differentially expressed genes within the QTL. The protein-coding gene with the highest expression difference was *Col4a6*, which exhibited a 9-fold reduction in A/J compared to C57BL/6J, consistent with decreased dopamine levels in A/J. Publicly available single cell RNA-seq data from developing human midbrain suggests that *Col4a6* is highly expressed in radial glia-like cells and neuronal progenitors, indicating possible involvement in neurogenesis. Interestingly, the lowered dopamine levels were accompanied by reduced striatal axonal branching of striatal DAns in A/J compared to C57BL/6J. Because *Col4a6* is known to control axogenesis in non-mammal model organisms, we hypothesize that different dopamine levels in mouse dorsal striatum are due to differences in axogenesis induced by varying COL4A6 levels during neural development.

## Introduction

Dopamine (DA), one of the main neurotransmitters in mammalian brain, is involved in several important activities such as motor and cognitive function. Using the amino acid tyrosine as a building block, dopaminergic neurons (DAns) can synthesize DA with two enzymes: tyrosine hydroxylase (TH) and DOPA-decarboxylase (Nagatsu et al., 2019). Two important populations of DAns, with distinct activities, locate in substantia nigra (SN) and ventral tegmental area (VTA) in the ventral midbrain. The DAns in SN project mainly to dorsal striatum and control motor function, while the ones in VTA project either to nucleus accumbens and amygdala, controlling reward and emotion, or to the cognitive centres in cortex and hippocampus, governing cognition and memory (Hassan and Benarroch, 2015; Vogt Weisenhorn et al., 2016). Both DAns are at the centre of neuroscience study because of their involvement in neurological diseases, notably VTA DAn in neuropsychiatric diseases and SN DAn in Parkinson’s disease (PD).

Because of its prevalence and costs to society with an aging population, PD has received a lot of attention over the last decade. Studies brought to light the importance of environmental and genetic risk factors in the variability of PD (Del Rey et al., 2018; Gilgun-Sherki et al., 2004; van Rooden et al., 2011; Jankovic et al., 1990). The different age of onset, severity, and rate of progression of PD motor symptoms, as well as the variable response to dopamine replacement therapies are likely due to human genetic polymorphism (Kaplan et al., 2014; Kalinderi et al., 2011). Yet the underlying factors governing the outcome of these polymorphisms are mostly unknown. As genetic studies with standardized and controlled environment are difficult to handle in human, mouse models are commonly used to study genetic variations. Indeed, mouse shares similar brain architecture and 99% of genes with human and allows cost-effective and controlled studies (Nadeau and Auwerx, 2019). Recombinant inbred mouse strains constitute interesting models to identify candidate genes with a method of quantitative trait locus (QTL) mapping (Peters et al., 2007). Collaborative Cross (CC) strains are a resource of recombinant inbred strains derived from eight founder strains for complex trait analysis, capturing an abundance of genetic diversity (Churchill et al., 2004). Genetic variation is also associated with phenotypic differences in the dopaminergic circuitry and the associated behaviours. Differences in the DAn cell number as well as in DA levels and protein trafficking have been shown between different strains of mice (Baker et al., 1980; Vadász et al., 1987; Vadasz et al., 1998; Zaborszky and Vadasz, 2001; Cabib et al., 2002). Motor behaviour and susceptibility to PD-toxin are also known to differ according to the mouse genetic background (Jong et al., 2010; Ingram et al., 1981; Brooks et al., 2004; Hamre et al., 1999). Altogether, previous works suggest that the dopaminergic circuitry is affected by genetic variation. But many factors underlying this variation are largely unknown.

In our study, we used CC mouse strains as a tool to map QTL related to the integrity of DAns. We measured dopamine level in the striata of eight CC founders and 32 derived CC strains, and observed that the striatal dopamine level was influenced by the genetic background of the strains. We identified a QTL associated with striatal dopamine level on chromosome X that contained 393 genes. The transcriptomic analysis of C57BL/6J and A/J ventral midbrains, two CC founders having large striatal dopamine level differences, revealed 24 differentially expressed genes within the QTL, with *Col4a6* showing the highest expression difference. Published studies using single cell RNA-seq data of developing human midbrain, have revealed a developmental expression profile for *Col4a6* indicating a role in neurogenesis (La Manno et al., 2016), and morphological studies in non-mammal models (Drosophila and zebrafish) indicate a role in axon guidance and neurite outgrowth (Mirre et al., 1992; Takeuchi et al., 2015). Consistently, measurements of TH-positive axons in projection areas of SN DAns (dorsal striatum) and of VTA DAns (piriform cortex, amygdala) revealed that axonal branching of SN DAns, but not that of VTA DAns, differed between C57BL/6J and A/J. However, the number of TH-positive neurons in the SN did not differ between these 2 strains. These observations indicate that differences in *Col4a6* expression lead to differences in dopaminergic striatal innervation.

## Materials and methods

### Animals

Eight parental founder strains (A/J, C57BL/6J, 129S1Sv/ImJ, CAST/EiJ, PWK/PhJ, WSB/EiJ, NOD/ShiLtJ, NZO/H1LtJ) and 32 CC strains (Supplementary Table S1), originally obtained from the University of North Carolina, Chapel Hill (UNC), were bred at Chapel Hill or at the Central Animal Facilities of the Helmholtz Centre for Infection Research (Braunschweig, Germany). 10 to 12 mice per group (mixed males and females) were anesthetized with a ketamine-medetomidine mix (150 and 1 mg/kg, respectively). Intracardiac perfusion was performed (phosphate-buffered saline) for each animal before dissecting the striatum and midbrain, immediately snap-frozen. The second hemibrain was fixed in paraformaldehyde (PFA) 4%. The experiments were performed according to the national guidelines of the animal welfare law in Germany (BGBl. I S. 1206, 1313 and BGBl. I S. 1934) and the European Communities Council Directive 2010/63/EU. The protocol was reviewed and approved by the ‘Niedersächsisches Landesamt für Verbraucherschutz und Lebensmittelsicherheit, Oldenburg, Germany’ (Permit Numbers: 33.9-42502-05-11A193, 33.19-42502-05-19A394), respecting the 3 Rs’ requirements for Animal Welfare.

### Dopamine measurements by gas chromatography-mass spectrometry (GC-MS)

Striatal DA was measured in 3-month-old CC founders and 32 CC strains. As we used two different methods to extract the metabolites from the tissues, we present the results as percentage of C57BL/6J. The tissue homogenization and metabolite extraction were performed at 4°C or lower to prevent changes in the metabolic profile.

The first method was described by Jaeger *et al*., 2015 (Jaeger et al., 2015). The striatum of each mouse was pulverized in a bead mill with grinding beads (7 mm). The samples were then homogenized in the bead mill with smaller grinding beads (1 mm) and the extraction fluid (methanol/distilled water, 40:8.5 v/v). The metabolites were extracted using a liquid-liquid extraction method first by addition of chloroform to the tissue fluid followed by distilled water. After shaking for 20 minutes at 1300 rpm at 4°C, the mixture was centrifuged for 5 minutes at 5000 x g at 4°C. The upper phase containing the polar metabolites was transferred to a sample vial for speed vacuum evaporation.

For both methods, the resulting dried samples were derivatized in an established procedure. 20 μL of pyridine (containing 20 mg/mL of methoxyamine hydrochloride) were added to the samples and incubated at 45 °C with continuous shaking for 90 min. Then 20 μL of MSTFA were added to the sample vial and incubated 30 min at 45 °C with continuous shaking.

After derivatization, the GC-MS analysis was performed with an Agilent 7890A GC, or 7890B for the second method, coupled to an Agilent 5975C inert XL mass selective detector (MSD) or 5977A for the second method (Diegem, Belgium). 1 μL of sample was injected into a Split/Splitless inlet operating in split mode (10:1) at 270 °C. Helium was used as a carrier gas with a constant flow rate of 1.2 mL/min. The second method was slightly modified to further reduce the runtime. The GC oven temperature was held 1 min at 80 °C (0.6 min at 90 °C for the second method) and increased at 36 °C/min to 260 °C (at 25 °C/min to 200 °C and held for 6 min for the second method). Then the temperature was increased at 22 °C/min and maintained at a constant temperature of 325 °C for 2 min (4 min post run time at 325 °C for the second method). The transfer line temperature was set constantly to 280 °C and the MSD was operating under electron ionization at 70 eV. As described by Jäger *et al*., 2016 (Jäger et al., 2016), a multi-analyte detection using a quadrupole analyzer in selected ion monitoring mode was used for a sensitive and precise quantification of DA and the internal standard DA-*d4*.

Statistical analysis was performed using the GraphPad Prism 8 software. After applying the Shapiro-Wilk test to assess the normality of our data, a one-way ANOVA was applied to analyse the striatal dopamine levels.

### Immunofluorescence

TH protein was measured by immunofluorescence in the dorsal striatum, amygdala, piriform cortex, and SN of 3-month-old C57BL/6J and A/J. From 6 to 14 hemibrains per group were fixed (PFA 4%) for 48h and stored in PBS with 0.2% of sodium azide. Parasagittal free floating sections (50 μm) were generated using a vibratome (Leica; VT 1000S) collected every 4^th^ sections in a tube containing a cryoprotective medium (polyvinyl pyrrolidone 1% w/v in PBS/ethylene glycol 1:1) and stored at −20°C. The lateral sections were collected for the striatum, amygdala and piriform cortex measurements, and the medial sections were collected for the SN measurements.

The sections were washed in PBS with 0.1% Triton X-100 (PBST) and permeabilized in PBS with 3% H_2_0_2_ and1.5% Triton X-100. The sections were then blocked for 1h in PBST with 5% of Bovine Serum Albumin (BSA) and incubated overnight with rabbit anti-TH antibody (1:1000, Millipore, AB152) diluted in PBST with 2% of BSA. After several washing, the sections were incubated for 2 hours with the secondary antibody (Alexa fluor™ 488 goat anti-rabbit 1:1000, Invitrogen), mounted on slides and embedded in fluoromount.

Imaging was performed using a Zeiss AxioImager Z1 upright microscope, coupled to a “Colibri” LED system, and an Mrm3 digital camera for image capture using the software Zeiss Zen 2 Blue. For each striatum, amygdala and piriform cortex section, three images of each brain area were taken at 40x magnification using the apotome system. After thresholding, the area occupied by TH stainings in each picture was determined using the FIJI imaging software (Schindelin et al., 2012; Masliah et al., 2000). For the SN sections, the pictures were taken at 10x magnification. The area occupied by TH positive neurons was measured in the region of interest corresponding to the SN using ImageJ FIJI software. We can distinguish four different areas of the SN. Each area was quantified and averaged separately and summed as a cumulated surface (mm^2^) (Ashrafi et al., 2017).

GraphPad Prism 8 software was used for the statistical analysis. After applying the Shapiro-Wilk test to assess the normality of our data, an unpaired t-test was applied to analyse the TH measurement in different areas.

### Quantitative trait locus mapping

The QTL mapping was done with http://gn2.genenetwork.org/. The dataset containing dopamine measurements of dorsal striata of 32 CC strains were located with search terms (Species: Mouse (mm10); Group: CC Family; Type: Phenotypes; Dataset: CC Phenotypes) and navigated to Record CCF_10001 and CCF_10002. The QTL mapping was done with GEMMA on all chromosomes, MAF >= 0.05 with LOCO method. The genome wide significance of QTL mapping on male and female are set by 500 permutation simulations with FDR under 5% for each scan.

### Estimation of ventral dopamine level heritability in CC strains

The broad sense heritability is estimated based on (Belknap, 1998). Briefly, the total phenotypic variance (Vp) is calculated on all CC strains. The genetic variance (Va) is estimated by the mean of within-strain variance. The heritability (H^2^) is calculated as Va/Vp.

## Results

### Differences in striatal dopamine levels across Collaborative Cross mice are under genetic control

To determine whether the reported phenotypic differences between CC mouse strains (Schoenrock et al., 2018; Schoenrock et al., 2020) are accompanied by differences in striatal DA levels, we measured DA levels in isolated dorsal striata from the eight inbred founder CC strains (Supplementary Table S1). In total, dopamine from 102 mice at the age of 3 months was measured by GC-MS. DA levels varied significantly across the founders (one-way ANOVA, p=0.0004, F=4.507), indicating strain-specific differences in DA levels in the nigrostriatal dopaminergic circuit (Figure 1). PWK/PhJ, A/J, and NOD/LtJ strains showed the lowest levels of DA, while the highest levels were detected in NZO/HILtJ, CAST/EiJ, and C57BL/6J mice. Thus, striatal DA levels appear to be under genetic control.

**Figure 1.**
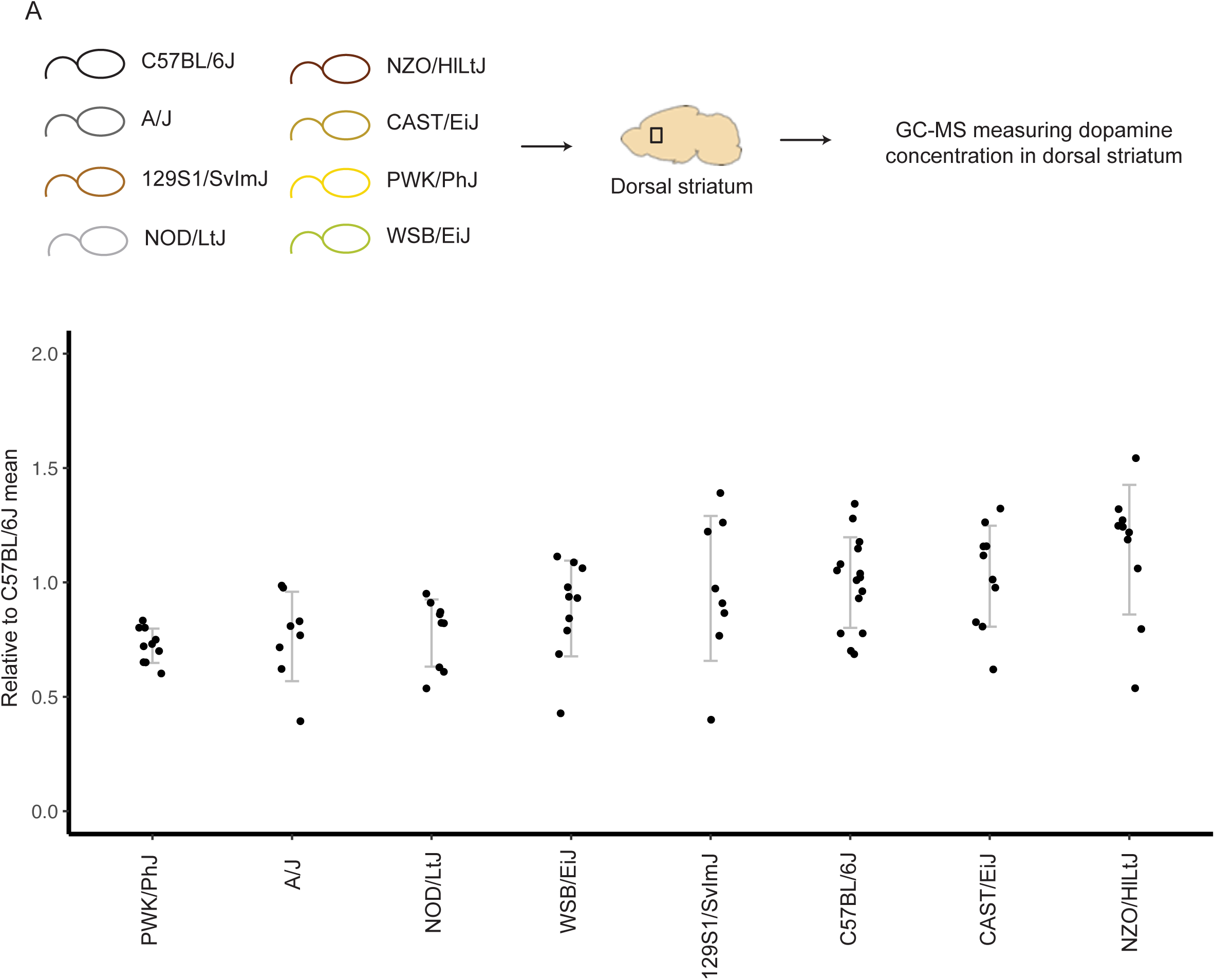
Striatal dopamine levels measured in the different Collaborative Cross (CC) founders. (A) Schematic representation of the experimental set-up. (B) Level of dopamine measured by GC-MS in the striatum of the different CC founders, expressed relative to the dopamine level of the common C57BL/6J strain. Data are expressed in mean ± standard deviation and the significance of differences was tested with one-way ANOVA (p=0.0004, F=4.507).

To investigate if variation in DA level is indeed inheritable, we measured the striatal DA level across 32 strains of CC mice. In total, we analysed 327 CC mice with similar number of mice from both sexes. The CC strains showed considerable variation in DA levels with a range of around 10 pmol/mg in both sexes (Figure 2). From these values, the estimated broad-sense heritability (H^2^) was calculated to be 0.52, indicating the DA level differences are inheritable, and associated genetic variation could be detected by QTL mapping. (Hegmann and Possidente, 1981) (see Methods for details).

**Figure 2.**
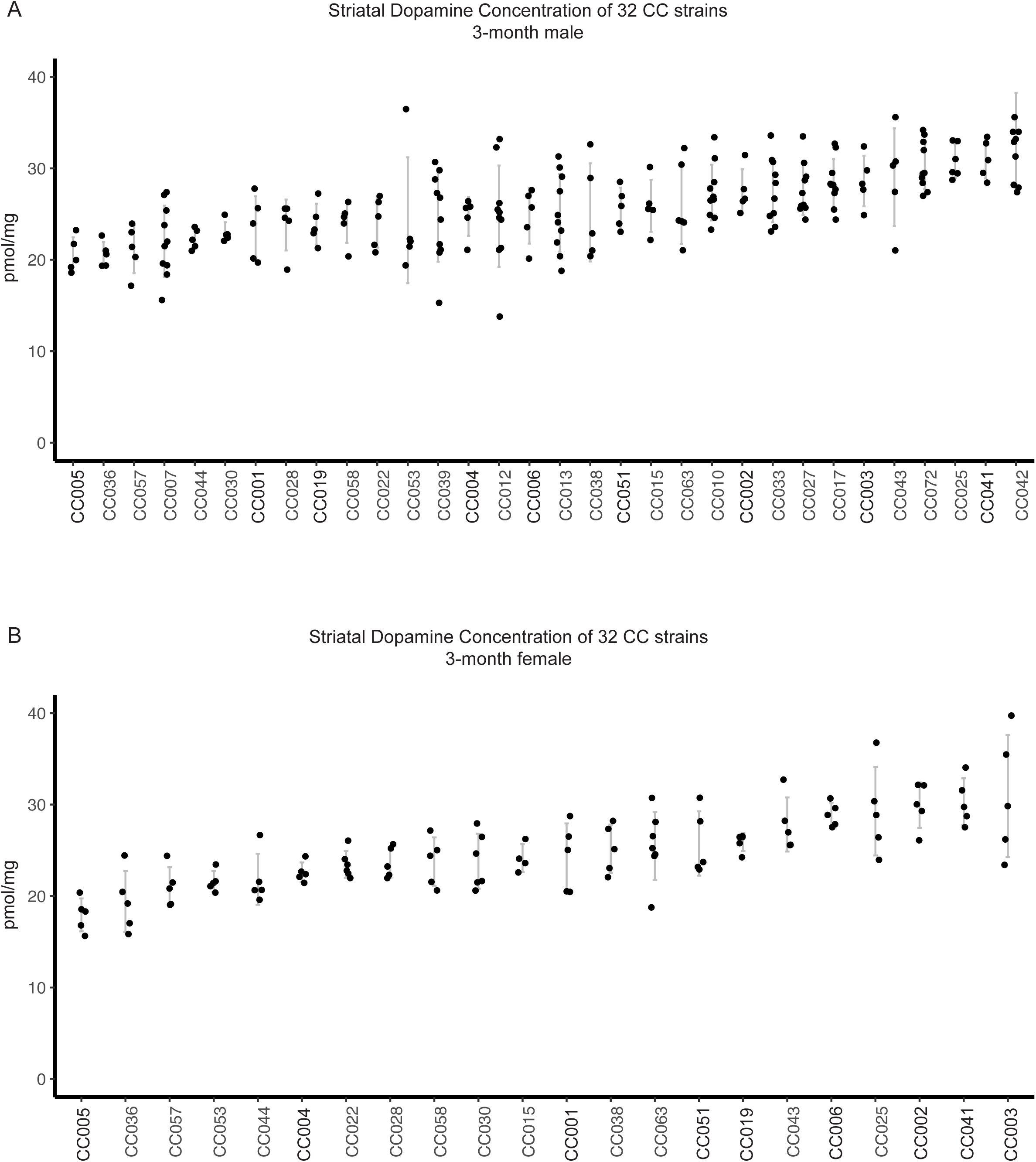
Striatal dopamine levels measured in the different collaborative cross (CC) strains. Levels of dopamine measured by GC-MS in the striatum of the different CC strains, expressed in pg/mg of tissue (mean ± standard deviation) and the significance of differences was tested with one-way ANOVA. (A) Level of dopamine in the striatum of CC males (p<0.0001, F=3.964). (B) Level of dopamine in the striatum of CC females (p<0.0001, F=7.435).

### QTL mapping associates a genomic locus on chromosome X with striatal dopamine levels

Identifying novel genetic regulators associated with striatal DA levels could help better understand the development of dopaminergic circuits and susceptibility to diseases, like PD. Therefore, to leverage the power of CC strains to identify trait-associated genetic loci at a good resolution, we performed QTL mapping based on the measured DA levels across the 32 CC strains. The mapping was performed separately for males and females, and the genome-wide significance results for all genetic markers are presented in Figure 3 and Supplementary Table S2. QTL mapping using the female data identified one genetic marker located on chromosome X at position 144.300241 Mb to be associated with DA levels, when applying the 95^th^ percentile threshold for genome-wide significance (-log_10_ p-value = 5.23) (Figure 3A). However, additional adjacent markers at downstream positions of 157.823410 Mb and 158.259643 Mb were also highly associated in females (-log_10_ p-value = 4.96 for both markers), while an upstream marker at position 136.176403 Mb showed strong association in males (-log10 p-value = 4.91). Taken together, our QTL analysis identifies of a combined region spanning over 32 Mb with high association to striatal DA levels on chromosome X.

**Figure 3.**
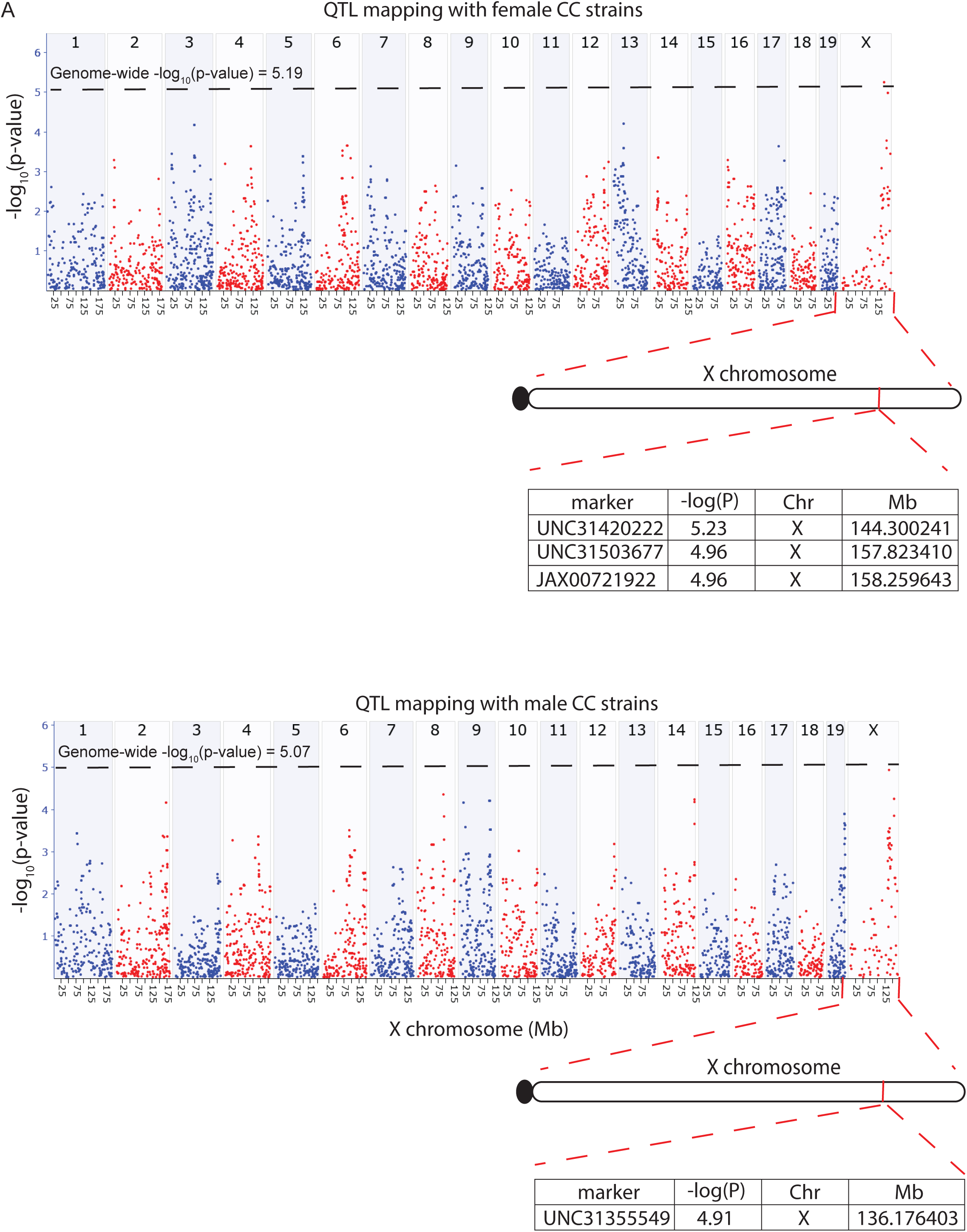
QTL mapping for dorsal striatum dopamine levels in CC strains. Plots show - log_10_ p-values (y-axis) of genetic markers across chromosome locations (x-axis). Horizontal dashed lines represent the 95^th^ percentile thresholds for genome-wide significance. (A) QTL mapping with female CC strains yielded the most significant QTL at chromosome X 144.3 Mb with -log(P) 5.23. (B) QTL mapping with male CC strains yielded the most significant QTL at chromosome X 136.2 Mb with -log(P) 4.91.

### *Col4a6* is a developmental gene with altered expression between mouse strains

The identified locus from position 131 Mb to 163 Mb on chromosome X includes 393 genes that could potentially be underlying the association with striatal DA levels. However, the vast majority (>95%) of trait-associated genetic variants are located outside of protein-coding genomic regions (Maurano et al., 2012). Recent advances in functional genomics analysis have revealed these non-coding variants to be highly enriched in gene regulatory regions such as enhancers where they can disrupt transcription factor binding and alter the target gene expression. Therefore, we asked whether such *cis*-regulatory variants could be affecting gene expression at our locus of interest in the SN of the midbrain, from where the DA neurons project to the dorsal striatum. To this aim, we took advantage of our recent transcriptomic profiling of ventral midbrains from C57BL/6J and A/J mice (Gui et al., 2020), two CC founder strains with significantly different levels of striatal DA (Figure 1, p=0.026, unpaired t-test). These two strains are also widely used laboratory mouse strains and have one of the largest difference in striatal dopamine (see above). Using the RNA-seq data we plotted the absolute log_2_-fold change for all 393 genes between C57BL/6J and A/J to identify those with altered gene expression (Figure 4A). Interestingly, only 24 genes showed significant differential gene expression (Supplementary Table S3). By far the largest fold change of all protein-coding genes was found for *Collagen 4a6* (*Col4a6*) gene, which showed over 9-fold difference between the 2 strains. The transcriptomic analysis was based on a total 24 mice, 12 from each strain and sex, with A/J displaying a significant reduction for *Col4a6* in both females and males compared to age-matched C57BL/6J (Figure 4B), consistent with a lower level of DA in the striatum of A/J (Figure 1).

**Figure 4.**
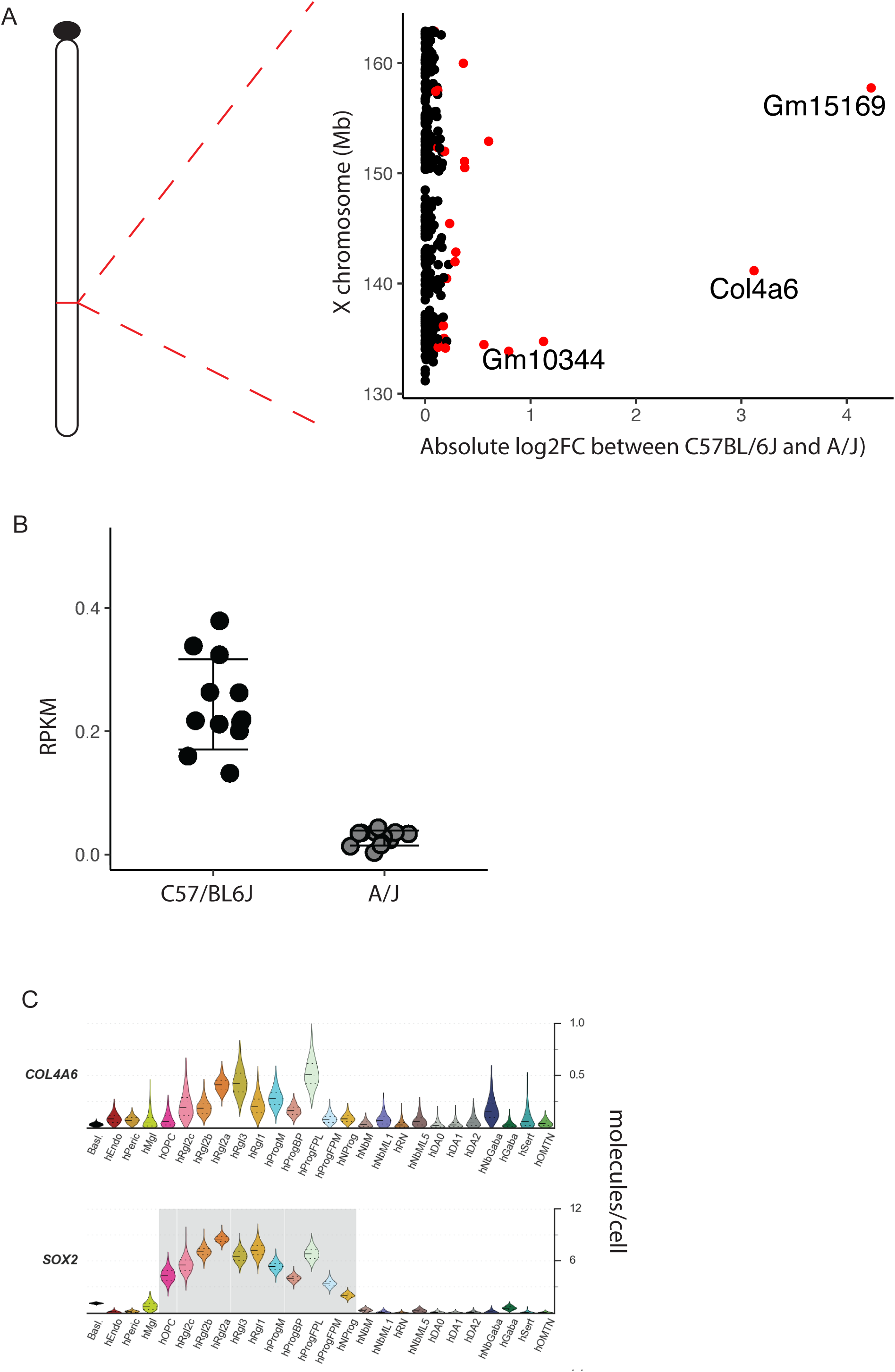
Col4a6 is the gene with highest differential expression between C57BL/6J and A/J at the identified chromosome X QTL. (A) Log2-fold changes of genes in chromosome X locus between 131Mb and 163 MB. Genes with adjusted p-value below 0.05 are labelled in red. (B) The expression (RPKM) of Col4a6 in the ventral midbrain of C57BL/6J and A/J. (C) The expression of Col4a6 and Sox2 during human midbrain during development based on scRNA-seq data from La Manno et al (La Manno et al., 2016). The two genes show similar expression profiles. Cell types are named with “h” to indicate human: Endo, endothelial cells; Peric, pericytes; Mgl, microglia; OPC, oligodendrocyte precursor cells; Rgl1-3, radial glia-like cells; NProg, neuronal progenitor; Prog, progenitor medial floorplate (FPM), lateral floorplate (FPL), midline (M), basal plate (BP); NbM, medial neuroblast; NbML1&5, mediolateral neuroblasts; RN, red nucleus; DA0-2, dopaminergic neurons; Gaba, GABAergic neurons; Sert, serotonergic; OMTN, oculomotor and trochlear nucleus.

While *Col4a6* expression was significantly lower in A/J, the overall abundance of expression in the adult C57BL/6J midbrain was also very low (<0.3 RPKM), indicating that, in adult mice, its expression is limited to only one or a few cell types, most likely endothelial cells, which have been reported to produce collagens (Gelse et al., 2003; Ricard-Blum, 2011). Collagen IV is an essential and abundant component of the basement membrane (Mao et al., 2015). In the nervous system, its function has been associated with axon guidance and neurite outgrowth in non-mammal model systems (Takeuchi et al., 2015; Mirre et al., 1992), and in cultured sympathetic neurons (Firla, 1990). The two angles of collagen IV function in the nervous system (regulation of neurogenesis and of neurite outgrowth and guidance) are probably intertwined.

To get a better idea about the cellular source of *COL4A6* during development, we observed its expression in published single cell RNA-seq (scRNA-seq) data corresponding to 26 cell types of the developing human midbrain (La Manno et al., 2016). *COL4A6* expression was highest in floor plate progenitors and selected subtypes radial glia-like cells, with very low expression detected in other cell types (Figure 4C). The expression profile of *COL4A6* closely followed the expression of *SOX2*, a key regulator of neurogenesis (Ferri et al., 2004). Consistently, previous screens for primary SOX2 target genes have found *COL4A6* expression to depend on SOX2 (Fang et al., 2011; Berezovsky et al., 2014).

Taken together, the expression profile of *COL4A6* implicates it as a developmental gene and its dependence on SOX2 indicates a possible role in neurogenesis. Indeed, previous work has shown that the zebrafish orthologs of type IV collagens, *col4a5* and *col4a6*, can control proper axonal guidance during zebrafish development (Takeuchi et al., 2015).

### Differences in striatal axonal branching between C57BL/6J and A/J mice

Based on the observed differences in striatal DA levels, the localization of *Col4a6* gene in the associated QTL, the distinct neurodevelopmental expression profile of *COL4A6*, and the previously described role of collagen IV in neurite outgrowth and guidance (see above), we hypothesized that DAn axonal fiber density could be altered specifically in the striatum of the mouse strains. To test this, we performed TH immunostaining on brain sections from the two founder strains C57BL/6J and A/J. The percentage of area occupied by TH in the DAn projection areas (dorsal striatum for SN and piriform cortex and amyglada for VTA) was used to compare axonal fiber density between the strains. Interestingly, 3-month-old A/J mice showed 29% lower TH fiber density in dorsal striatum compared to C57BL/6J (unpaired t-test, p=0.0059), consistent with lower DA levels in A/J (Figure 5). No such differences could be observed in amygdala or in piriform cortex, suggesting that the differences observed between the two mouse strains appear to be specific to the dorsal striatum.

**Figure 5.**
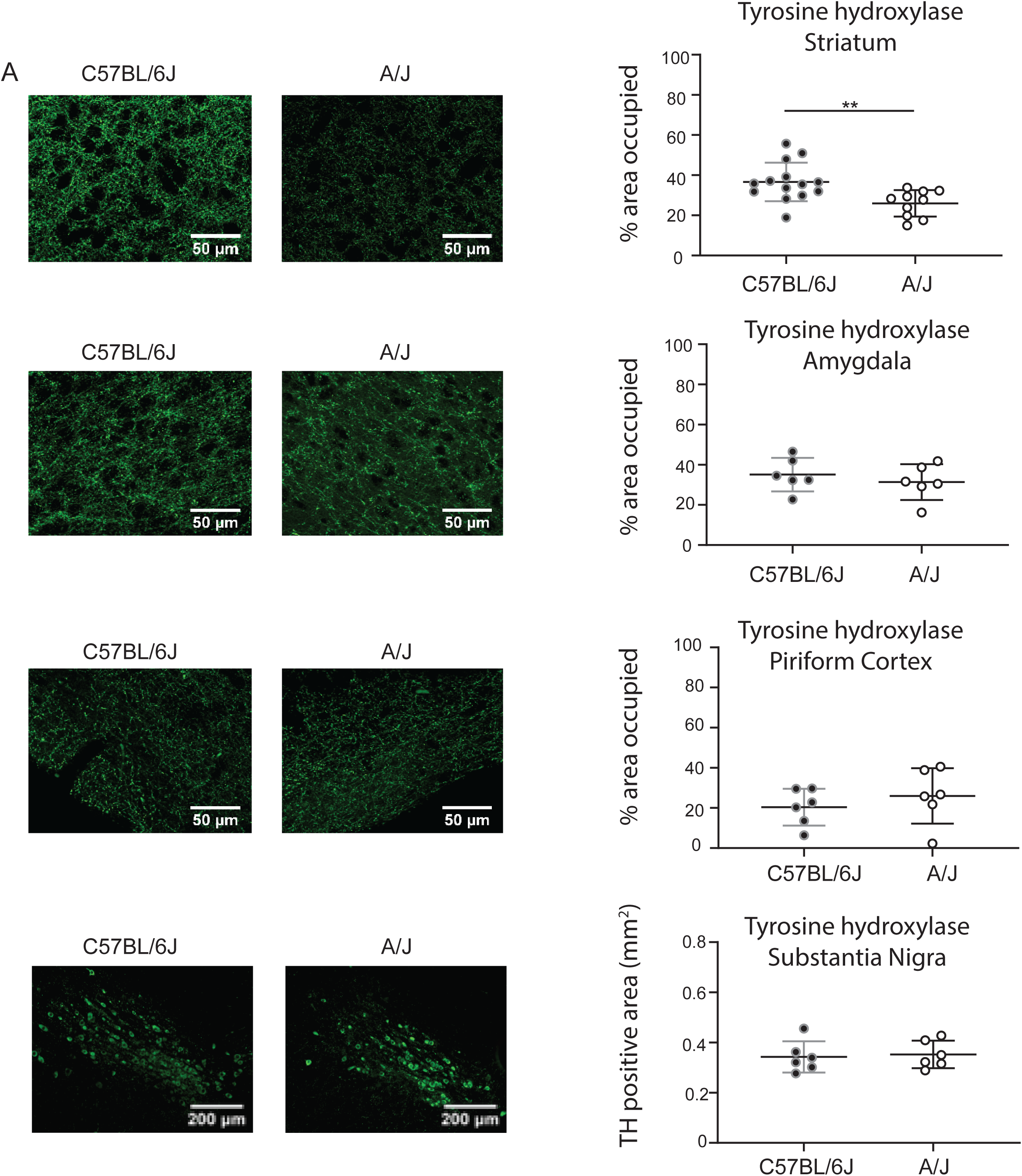
Measure of tyrosine hydroxylase (TH) by immunofluorescence in the brain of 3-month-old C57BL/6J and A/J. (A, B) TH in striatum. (C, D) TH in amygdala. (E, F) TH in piriform cortex. (G, H) TH in substantia nigra. (A, C, E) Images of TH stainings in striatum, amygdala and piriform cortex, magnification 40X. Scales bar: 50 µm (G) Images of TH stainings in substantia nigra, magnification 10X. Scale bar: 200 µm (B, D, F) Quantification of the percentage area occupied by TH from stainings in striatum, amygdala and piriform cortex. (H) Quantification of the TH-positive area in substantia nigra. Data are expressed in mean ± standard deviation and were analysed with unpaired t-test (p=0.0059 in the striatum (**), p=0.47 in amygdala, p=0.43 in piriform cortex and p=0.78 in SN).

To determine if the number of DAn differed between C57BL/6J and A/J mice, we estimated the amount of these neurons in the 2 strains as described in Materials and Methods. We observed no difference in the number of TH-positive neurons between the two strains, implying that differences observed in striatal TH positive axons reflect a difference in branching of DAn, rather than their number of the SN (Figure 5). Hence, our data points to a role of COL4A6 in modulation of the branching of DAn of the SN, but not that of DAn of the VTA.

Taken together, our QTL analysis allowed us to identify *Col4a6* as a new gene putatively involved in the DAns neurogenesis and axonal branching in the dorsal striatum.

## Discussion

While the DAns residing in the SN are of central interest to translational neuroscience research because of their unique properties that renders them susceptible (Surmeier, 2018), a lot of the mechanisms surrounding their basic properties remain unknown.

In this study, we used CC strains to identify QTL and new candidate genes regulating the integrity of the nigrostriatal circuitry. The variations of striatal dopamine levels between CC strains demonstrate an inheritable part of this trait, which could also underlie the variability in human populations. Together with previous transcriptomic data in the midbrain of two founder strains, our results point to *Col4a6* playing an important role in neurogenesis and axonal branching of striatal DAns.

Despite the success of human GWAS methodologies to decipher phenotype-genotype associations, human tools lack proper standardized and controlled conditions. To overcome these limitations, CC mouse strains were generated to provide a model for heterogeneous human population (Churchill et al., 2004). Genetic diversity of CC mouse model provides more precise QTL mapping results than conventional mapping populations. The wide phenotypic range of around 10 pmol/mg DA enabled us to map a significant QTL of about 32 Mb with a reasonable number of 32 CC strains and eight founder strains. Our previous transcriptomic analysis coming from two CC founders with significantly different levels of striatal DA (Gui et al., 2020) provide useful data to narrow the QTL on chromosome X to 24 DEG, *Col4a6* showing a 9-fold difference in C57BL/6J compared to A/J, two of the CC founder strains that had one of the largest striatal DA differences. To find out more about the role of Col4a6 in this brain region, we started to look at data available from the developing human midbrain (La Manno et al., 2016), which suggests an important role of *Col4a6* in the DAns neurogenesis. Collagen IV alpha-6 chain is one of the six subunits of type IV collagen, a major component of basement membranes. Collagen IV is a member of the collagen family of glycoproteins, which themselves are constituents of the extracellular matrix (ECM), and among the most abundant proteins in the animal kingdom (Vecino and Kwok, 2016). ECM proteins, in particular collagens, in the nervous system play key roles in development, in cellular maintenance and repair, and in tissue responses to diseases involving injury or neoplasia (Rutka et al., 1988). Collagens in the PNS provide a scaffold for Schwann cells and support neurite outgrowth (Lein et al., 1991; Chen et al., 2015). As stated above, a central role for collagen IV in axon guidance and neurite outgrowth is also supported by studies in simple model organisms (Mirre et al., 1992; Takeuchi et al., 2015). Collagens, including type IV, in the nervous system are produced primarily by cells of mesodermal origin, the endothelial cells, and are found, together with other forms ECM proteins, in the basement membrane of cerebral blood vessels and at the glia limitans (Rutka et al., 1988). Upon injury, collagen expression and secretion together with that of other ECM proteins, by glial cells appears (Liesi and Kauppila, 2002). While evidence suggest that, in some scenarios, this process is supportive of neurite outgrowth (see above), in rats, by contributing to the formation of the glial scar, it is thought to inhibit axonal regeneration (Liesi and Kauppila, 2002). Interestingly though, engineered biopolymer scaffolds containing collagen are being explored as therapeutic support for nerve repair after injury (Li and Dai, 2018). Thus, the role of collagens in neuronal maintenance and repair may depend on a complex balance of opposing actions.

Our study shows for the first time that one subunit of collagen IV, subunit 6, is a candidate for regulating axonal branching in the CNS of a mammal. Because we had already observed large striatal DA differences and large midbrain *Col4a6* expression differences in C56BL/^J versus A/J mice, we then tested if these two mouse strains also showed differences in striatal axonal branching. We therefore measured TH-positive axons in the dorsal striatum C57BL/6J and A/J by quantitative immunofluorescence. We observed less TH-positive axons in the dorsal striatum of A/J compared to C57BL/6J mice. The determine if this was due to a difference in the number of TH positive neurons in the SN between these two strains, we also estimated those numbers, but did not find a difference. Thus, the lower TH-positive axons we observe in the striata of A/J mice most likely reflect a lesser axonal branching of the DAns.

To ensure that our strain differences affect the nigrostriatal circuitry and not all projecting DAns of the midbrain, we assessed brain areas that receive projections from the VTA. Based on our results from the amygdala and piriform cortex, phenotypic differences observed in the striata of C57BL/6J and A/J are not present in the other midbrain dopaminergic circuits. Other studies indicate that more factors regulating nigrostriatal integrity remain to be found. Studying mouse strains other than those used in our study, Baker *et al*., 1980 (Baker et al., 1980) and Zaborsky *et al*., 2001 (Zaborszky and Vadasz, 2001) reported strain-dependent differences in the number of midbrain populations of DAns. On a functional level, other studies found strain-specific differences in motor behaviour, such as lower motor activity, balance and exploratory skills displayed by A/J compared to C57BL/6J (Jong et al., 2010). Finally, recent studies showed significant effect of the genetic background on locomotor behaviour (Schoenrock et al., 2020) and response to cocaine (Schoenrock et al., 2018) using recombinant inbred intercrosses generated from CC strains, illustrating how the CC strains can serve as useful model for identifying further QTLs and genetic variants that govern structure and function of DAn’s.

In this study, using a combination of biochemical and neuropathological analyses combined with QTL mapping in the CC mouse population, we highlight *Col4a6* as a new gene candidate regulating the axonal branching in the nigrostriatal dopaminergic system of mammals. Because these are the structures that are affected early in PD (Kordower et al., 2013), we propose that a better understanding of the actions of collagen IV on these neurons may open the way for novel neuroprotection therapies.

## Acknowledgements

We would like to thank Drs Aurélien Ginolhac and Anthoula Gaigneaux for their support with bioinformatic analysis and EMBL Gene Core at Heidelberg for support with high-throughput sequencing, and Dr Djalil Coowar (Animal Facility of University of Luxembourg) for help with breeding of experimental mice. KS would like to thank the animal caretakers at the Central Animal Facilities of the HZI for maintaining the mice. The computational analysis presented in this paper were carried out using the HPC facilities of the University of Luxembourg (68).

